# Point mutations in Arf1 reveal cooperative effects of the N-terminal region and myristate for GTPase-activating protein catalytic activity

**DOI:** 10.1101/2023.11.15.567322

**Authors:** Eric M. Rosenberg, Xiaoying Jian, Olivier Soubias, Rebekah A. Jackson, Erin Gladu, Emily Andersen, Lothar Esser, Alexander J. Sodt, Di Xia, R. Andrew Byrd, Paul A. Randazzo

**Affiliations:** Laboratory of Cellular and Molecular Biology, Center for Cancer Research, National Cancer Institute; Bethesda, MD, USA; Section of Macromolecular NMR, Center for Structural Biology Laboratory, Center for Cancer Research, National Cancer Institute; Frederick, MD, USA; Laboratory of Cell Biology, Center for Cancer Research, National Cancer Institute; Bethesda, Maryland, USA; Unit of Membrane Chemical Physics, Eunice Kennedy Shriver National Institute of Child Health and Human Development; Bethesda, MD, USA

## Abstract

The ADP-ribosylation factors (Arfs) constitute a family of small GTPases within the Ras superfamily, with a distinguishing structural feature of a hypervariable N-terminal extension of the G domain modified with myristate. Arf proteins, including Arf1, have roles in membrane trafficking and cytoskeletal dynamics. While screening for Arf1:small molecule co-crystals, we serendipitously solved the crystal structure of the non-myristoylated engineered mutation [L8K]Arf1 in complex with a GDP analogue. Like wild-type (WT) non-myristoylated Arf1•GDP, we observed that [L8K]Arf1 exhibited an N-terminal helix that occludes the hydrophobic cavity that is occupied by the myristoyl group in the GDP-bound state of the native protein. However, the helices were offset from one another due to the L8K mutation, with a significant change in position of the hinge region connecting the N-terminus to the G domain. Hypothesizing that the observed effects on behavior of the N-terminus affects interaction with regulatory proteins, we mutated two hydrophobic residues to examine the role of the N-terminal extension for interaction with guanine nucleotide exchange factors (GEFs) and GTPase Activating Proteins (GAPs). Different than previous studies, all mutations were examined in the context of myristoylated Arf. Mutations had little or no effect on spontaneous or GEF-catalyzed guanine nucleotide exchange but did affect interaction with GAPs. [F13A]myrArf1 was less than 1/2500, 1/1500, and 1/200 efficient as substrate for the GAPs ASAP1, ARAP1 and AGAP1; however, [L8A/F13A]myrArf1 was similar to WT myrArf1. We hypothesized that the myristate moiety associates with the N-terminal extension to alter its structure, thereby affecting its function. Using molecular dynamics simulations, the effect of the mutations on forming alpha helices was examined, yet no differences were detected. The results indicate that lipid modifications of GTPases and consequent anchoring to a membrane influences protein function beyond simple membrane localization. Hypothetical mechanisms are discussed.

## Introduction

The ADP-ribosylation factor (Arf) family of small GTPases within the Ras superfamily regulate membrane trafficking and cytoskeletal reorganization and are being investigated for roles in cancer progression (1, 2). Arf function depends on cycling between GDP- and GTP-bound states. Different than other families of GTPases, intrinsic exchange rates and GTPase rates of Arf proteins are slow or negligible, and they therefore depend on guanine nucleotide exchange factors (GEFs) and GTPase-activating proteins (GAPs) for activity (3, 4). The G domain architecture of Arf proteins is similar to other G proteins, consisting of a P loop, G4 motif, and G5 motif that bind to guanine nucleotides, as well as switch regions I and II (and interswitch region) that are conformationally different depending on the guanine nucleotide bound (5). Like other small GTPases, Arf proteins have a lipidated hypervariable region (HVR), but different than other small GTPases, the HVR is an N-terminal extension of typically 16 amino acids from the G domain that is co-translationally modified with myristate on the glycine at position 2 (6, 7). In cells, both the N-terminal region and the myristate are essential; [Δ17]Arf1 and [G2A]Arf1 have no detectable activity in yeast or mammalian cells and peptides comprised of the N-terminal regions of Arf proteins block Arf functions (8, 9). The possible coupling of the myristate with residues of the N-terminus is understudied and the molecular bases for the requirement of myristate and the N-terminal region are still being discovered.

The myristate is often considered to function as a membrane anchor, important for confining proteins to specific endomembranes but otherwise not critical for interaction with target proteins (10, 11). Prior to this report, results examining Arf GAPs appear to be consistent with this hypothesis. [Δ17]Arf1 is a poor substrate for Arf GAPs (12, 13). In the context of the full-length non-myristoylated protein, specific residues within the N-terminal region of Arf1 were found to be critical, including lysines at positions 10, 15, and 16 (13, 14). The possible interaction of the myristate with the N-terminal region in determining GAP activity was not pursued beyond an examination of wild-type (WT) Arf1, in which myristate did not affect GAP activity (15, 16). In the NMR structure of yeast myristoylated Arf•GTP, the myristate was buried in lipid bicelles where it formed significant contacts with the conserved residue leucine 8, a residue not critical for GAP activity (in the context of non-myristoylated protein) (14, 17). These results were interpreted as consistent with the conclusion that myristate does not have a role in GAP activity.

The effects of myristate and the N-terminal region on guanine nucleotide exchange factors are more complex, raising the possibility of cooperative interactions between the structures. Full-length, non-myristoylated Arf1 is a poorer substrate for GEFs than is [Δ17]Arf1, while full-length myristoylated Arf1 (hereby referred to as “myrArf”) is a better substrate than [Δ17]Arf1 (18, 19). The myristate has a clear functional role by itself: integrated structural biology approaches have revealed a role for the myristate in anchoring the transition complex of nucleotide exchange to a hydrophobic surface (20, 21). However, a possible cooperative function of the N-terminal region and the myristate was revealed by a comparison of the structures of myristoylated and non-myristoylated Arf•GDP. Whereas the N-terminal region forms an alpha helix that occupies a hydrophobic cavity in non-myristoylated Arf•GDP, the myristate occupies the same site in myrArf with a disordered N-terminal region floating on top, potentially increasing accessibility for the GEF (22, 23). Thus, based on these limited results, the myristate and N-terminal regions of Arf proteins might have codependent function.

Here, we solved the crystal structure of the non-myristoylated engineered Arf1 mutant, [L8K]Arf1, bound to a GDP analogue (G3D). This mutant was previously generated to remove the requirement for a membrane surface for GTP/GDP exchange (24). The differences in the structure of the N-terminus and its integration within the protein from that observed with WT Arf1 motivated us to reexamine the role of the N-terminus in nucleotide exchange and GAP-dependent GTPase activity in the context of myristoylated Arf. We focused on two hydrophobic residues in the N-terminus of Arf1, leucine 8 (L8) and phenylalanine 13 (F13), as there is evidence for contacts between L8 and the myristate and because F13 is able to contact the membrane in both Arf•GDP and Arf•GTP. Although mutating L8 and F13 had little or no effect on spontaneous or catalyzed nucleotide exchange, mutating F13 affected GAP-induced GTP hydrolysis by several orders of magnitude. The effect of mutating F13 was reversed by simultaneously mutating L8. In previous studies, mutations of the same residues in non-myristoylated Arf1 had little or no effect on GAP-induced GTP hydrolysis (14). Molecular dynamics (MD) simulations on myristoylated N-terminal peptides suggested that the mutations do not affect peptide secondary structure, indicating that there is yet another undefined mechanism by which these mutations affect Arf GAP activity. Together, our results reveal a cooperative function of the myristate and the N-terminus in GAP-dependent regulation of Arf, which might extrapolate to Arf effectors and other Ras superfamily members.

## Results

### The crystal structure of non-myristoylated [L8K]Arf1 in complex with a GDP analogue exhibits an unexpected N-terminal shift

While conducting crystal screens for small molecules that bind to Arf proteins (25), we serendipitously crystallized non-myristoylated [L8K]Arf1 in complex with a GDP analogue (namely guanosine-3’-monophosphate-5’-diphosphate or G3D, discussed below). These crystals formed regardless of the presence of the small molecule ligands, and we therefore solved the structure of an *apo* crystal to 1.75 Å in order to ensure that no features of the data were present as a result of binding to these small molecules. Crystallographic statistics are shown in Table 1, and the overall structure of [L8K]Arf1•G3D (PDB accession code 8SDW) is shown in Fig. 1A. There is only a single monomer of [L8K]Arf1 in the crystal asymmetric unit, with residues 6 – 180 visible in the structure; in addition to residues 2 – 5, part of switch II (residues 70 – 74) was too disordered to be modeled. The macromolecule is clearly bound to a magnesium ion as well as a guanine nucleotide that was initially thought to be GDP, but was later determined to be G3D, a GDP molecule with a phosphate in the 3’ position (Supplemental Fig. S1A). G3D has been found in other structures of small GTPases including Arf (21, 26-28), Arf-like (Arl) (29), and Rab (30) structures, as well as a presumed nucleoside kinase, yorR (31), although its significance in these structures has yet to be determined and is likely an artifact of overexpression in *E. coli*. Although outside of the scope of this work, G3D has a known role in the bacterial stringent response, which occurs when bacteria are under conditions of amino acid starvation during growth (32-34).

**Figure 1.**
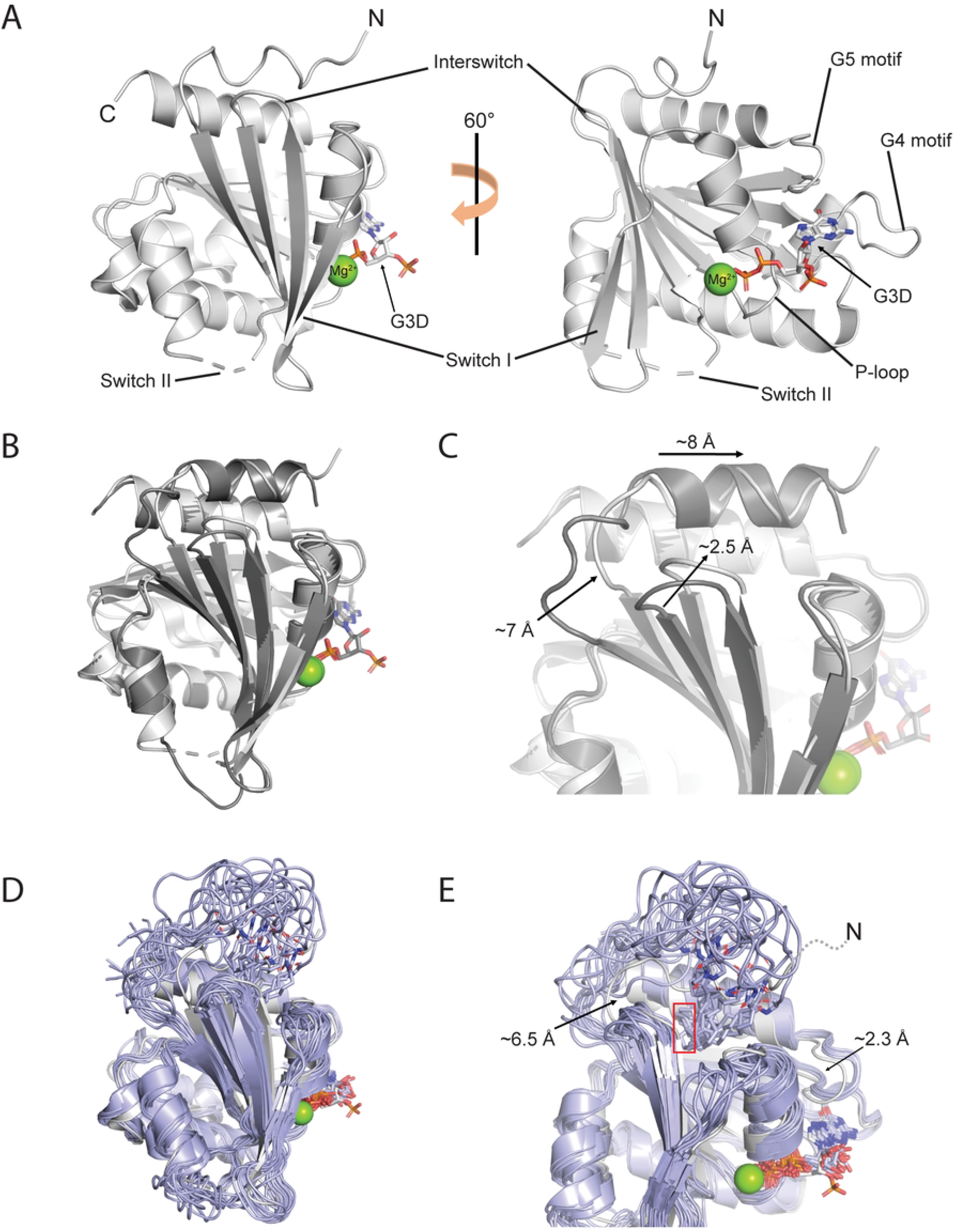
(**A**) Crystal structure of [L8K]Arf1 in complex with guanosine-3’-monophosphate-5’-diphosphate (G3D). N- and C-termini, G domain motifs, and magnesium ion are labeled. Part of switch II could not be modeled and is depicted with dashed lines. (**B**) Overall superimposition of [L8K]Arf1•G3D (light grey) and non-myristoylated WT Arf1•GDP crystal structures (dark grey, PDB: 1HUR (22)). (**C**) Structural differences between [L8K]Arf1•G3D and non-myristoylated WT Arf1•GDP. Movements of the N-terminal HVR, hinge region connecting the HVR and G domain, and interswitch region in the [L8K] structure compared to WT are depicted with arrows. Structures are colored as described in (B). (**D**) Overall superimposition of the [L8K]Arf1•G3D crystal structure (light grey) and the *S. cerevisiae* WT myrArf1•GDP NMR structure (light purple, PDB: 2K5U (23)). Note that all NMR states of the yeast myrArf1 structure are depicted. (**E**) Structural differences between [L8K]Arf1•G3D and yeast WT myrArf1•GDP. Movements of the hinge region connecting the HVR and G domain as well as G5 motif in the [L8K] structure compared to yeast myrArf1 are depicted with arrows. Differences between the N-termini are also highlighted: residues 2 – 5 in the [L8K] structure are disordered and could not be modeled, but their relative positioning as an extension of the modeled N-terminal residues are depicted with a dashed line; the positioning of the distal end of the myristoyl moiety in the yeast myrArf1 structure is shown with a red rectangle. Structures are colored as described in (D).

**Table 1.** [L8K]Arf1•G3D crystal data collection and refinement statistics.

|  | [L8K]Arf1•G3D (PDB: 8SDW) |
| --- | --- |
| **Data collection** |  |
| Space group | P2<sub>1</sub>2<sub>1</sub>2<sub>1</sub> |
| Cell dimensions |  |
| *a*, *b*, *c* (Å) | 41.65, 55.02, 78.47 |
| $\alpha$, $\beta$, $\gamma$ (°) | 90.0, 90.0, 90.0 |
| Resolution (Å) | 50.00 - 1.75 (1.78 – 1.75) |
| *R* p.i.m. | 0.025 (0.078) |
| CC<sub>1/2</sub> | 0.996 (0.976) |
| *I* / $\sigma I$ | 27.10 (9.33) |
| Completeness (%) | 98.7 (89.8) |
| Redundancy | 6.2 (5.8) |
| **Refinement** |  |
| Resolution (Å) | 39.24 - 1.75 (1.81 – 1.75) |
| No. reflections | 18523 |
| *R*<sub>work</sub> / *R*<sub>free</sub> | 0.19/0.21 |
| No. atoms | 1651 |
| Protein | 1322 |
| Ligand/ion | 33 |
| Water | 296 |
| *B*-factors |  |
| Protein | 20.4 |
| Ligand/ion | 17.1 |
| Water | 30.1 |
| R.m.s. deviations |  |
| Bond lengths (Å) | 0.005 |
| Bond angles (°) | 1.01 |

The data set was obtained from a single crystal, and the resulting data collection and refinement statistics are shown. Values in parentheses are for the highest resolution shell.

The crystal structure of [L8K]Arf1 is largely similar to that of full-length non-myristoylated WT Arf1 in complex with GDP (PDB accession code 1HUR (22)) with only minor structural rearrangements (Fig. 1B, 1C). The most striking difference between the WT and [L8K] structures is in the positioning of their N-terminal residues; while both structures exhibit amphipathic helices that occlude the hydrophobic cavity where the myristoyl group is typically found in the native protein (see next section), the helices are offset by approximately 8 Å, with the hinge region between the N-terminal helix and β1 being approximately 7 Å more extended in the WT structure than in the [L8K] structure (Fig. 1C). Calculating the RMSD of a monomer from 1HUR with the [L8K]Arf1 monomer highlights the large structural changes apparent as a result of the N-terminal shift. When comparing only residues 18 – 180, the RMSD values between the structures are 0.82 and 0.81 Å (between either α-carbons and backbone atoms); by including residues 6 – 17 in the analysis, the RMSD values are 1.95 and 1.98 Å. Another difference between the structures is the interswitch region, which in the [L8K] structure moves approximately 2.5 Å towards the cavity compared to the WT structure (Fig. 1C). However, while the changes in the N-terminal helix of [L8K]Arf1 do not appear to be a result of crystal contacts (nor can crystal contacts fully explain the positioning of the N-terminal helix in the WT structure (22), data not shown), the interswitch region is adjacent to a symmetry mate in the crystal lattice, and therefore the movement of the interswitch region may be an artifact of crystallization rather than a feature brought about the L8K mutation (Supplemental Fig. S1B). We also note that, along this same interface, a pair of lysine residues in the N-terminal region and interswitch region (K10 and K59) appear to form ionic interactions with the 3’ phosphate in G3D (Supplemental Fig. S1B).

Although the structure of human myrArf1•GTP has not been determined (Zhang *et al., Nat. Commun*, in press), the NMR structure of *S. cerevisiae* myrArf1•GDP is available (PDB accession code 2K5U (17)). *S. cerevisiae* and human Arf1 are the same length (181 residues) and share 77% identity/88% similarity, with the differences scattered throughout their primary structure. Their N-terminal regions are also the same length, namely 16 amino acids (residues 2 – 17) after processing. The yeast myrArf1 structure highlights the fact that the N-terminal region of Arf1 is typically disordered in the GDP-bound state, with the myristoyl moiety inserting into a hydrophobic cavity between the interswitch region and the C-terminal alpha helix (Fig. 1D, 1E). By comparing the yeast myrArf1•GDP and [L8K]Arf1•G3D structures, it is apparent that the most obvious differences lie in the positioning of their N-terminal regions; in addition, the hinge region between the N-terminal region and the first beta strand is approximately 6.5 Å more extended in the yeast myrArf1 structure than in the [L8K]Arf1 structure, similar to WT non-myrArf1 (Fig. 1C, 1E). [L8K]Arf1•G3D also shows a ∼2.3 Å inward movement of the G5 motif compared to yeast myrArf1•GDP (Fig. 1E). The inward movement of the G5 motif may be mediated by crystal contacts (Supplemental Fig. S1C). Alternatively, movement of the G5 motif may be a difference between yeast and human Arf1 as the G5 motif is the same position in the WT non-myrArf1 structure (Fig. 1B, 1C), or could even be driven the myristate itself, which is only present in the yeast but not human structures.

### The hydrophobic cavity of Arf1•GDP is occupied by either a myristoyl moiety in myrArf1 or the hydrophobic residues in the N-terminal region of non-myristoylated Arf1

Examination of the location of the L8K mutation revealed that this residue, while located inside the hydrophobic cavity in the WT structure as L8, is instead found outside of the hydrophobic cavity when mutated to lysine (Fig. 2). Interestingly, the N-terminal shift as a result of the L8K mutation is also achieved by redundancies in the N-terminal sequence of Arf1—while the hydrophobic cavity in the WT structure is filled by residues F5, L8, F9, L12, and partly M18, the cavity in the [L8K] structure is instead filled by F9, L12, F13, and M18 to a greater degree (Fig. 2A – 2C, Table 2). Indeed, the amino acid side chain positions of I4, F5, L8, and F9 are almost identical to the side chain positions of K8, F9, L12, and F13 in the [L8K] structure (Fig. 2A, Table 2). L12 in the WT structure is the only residue in the cavity that is not replaced by an identical amino acid in the [L8K] structure; instead, M18 shifts over by more than 3 Å in order to fill in the cavity (Fig. 2A, Table 2). Similarly, the movement of the interswitch region in the [L8K] structure shrinks the hydrophobic cavity by approximately 10 – 25% compared to the WT structure, potentially minimizing the free energy of the protein. The ordered positions of these N-terminal residues are in stark contrast to their positions in yeast myrArf1•GDP, in which they are exposed to solvent and only the myristate moiety occupies the hydrophobic cavity (Fig. 2D).

**Figure 2.**
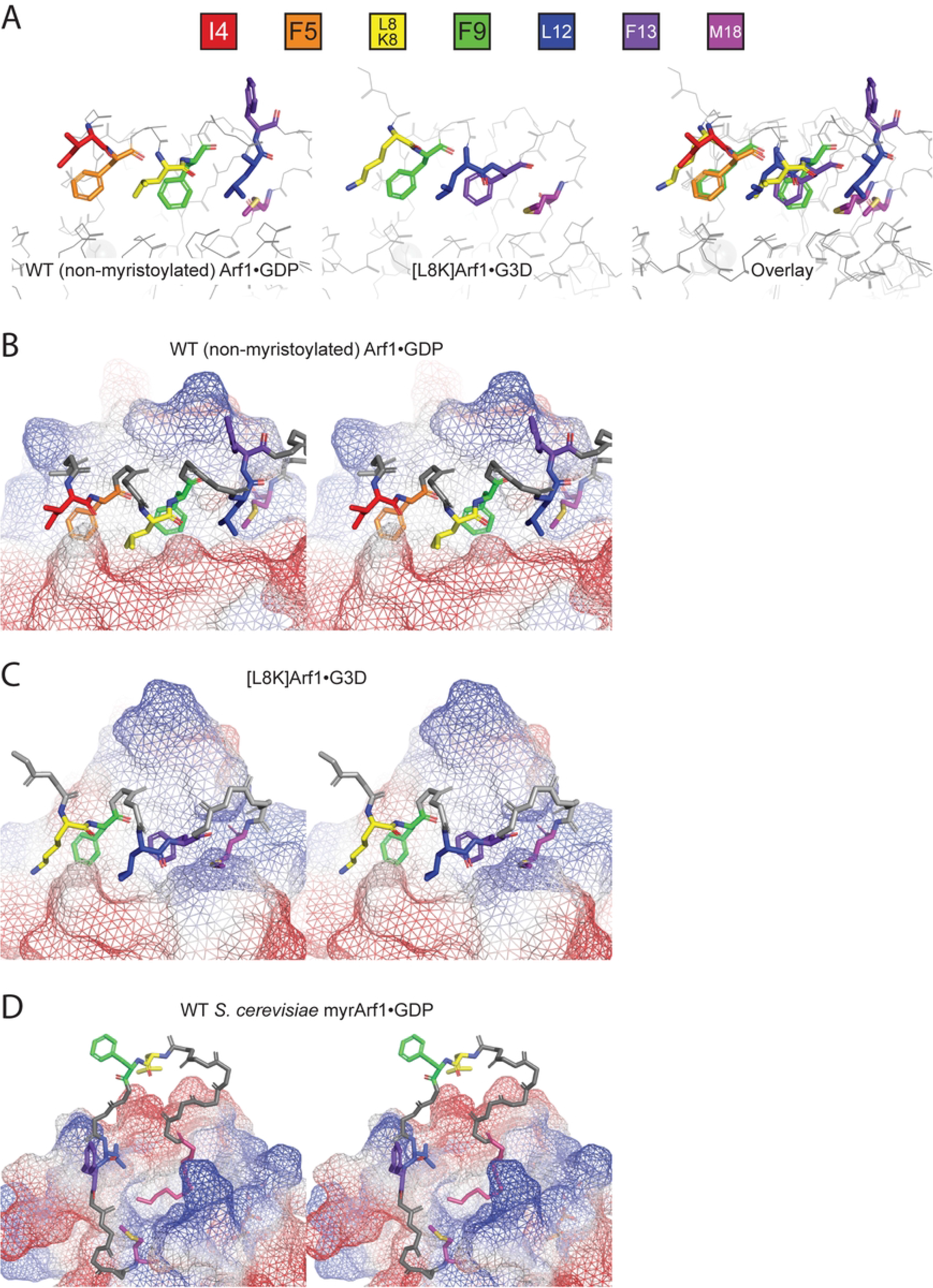
(**A**) Comparison of the positions of residues within the N-terminal extensions of non-myristoylated WT Arf1•GDP (left) and [L8K]Arf1•G3D (middle). An overlay of the two structures is shown on the right. For both structures, the amino acids that fill the hydrophobic cavity in the G domain are shown as colored sticks (key on top). (**B**) Stereoscopic view of the non-myristoylated WT Arf1•GDP N-terminal extension. The surface of the G domain is depicted in a mesh, with blue and red indicating positive and negative surface charges. Amino acids that fill the hydrophobic cavity are colored as described in (A). (**C**) Stereoscopic view of the [L8K]Arf1•G3D N-terminal extension. Colors of amino acids are as described in (A), and other features as described in (B). (**D**) Stereoscopic view of a single pose of the *S. cerevisiae* WT myrArf1•GDP N-terminal extension. Note that this view is from a different angle than those shown in (B) and (C), for clarity. The myristoyl moiety is shown in pink, and colors of amino acids that are conserved with human Arf1 (i.e., L8, F9, F13, and M18, see Fig. 3C) are as described in (A). Other features are as described in (B).

**Table 2.**
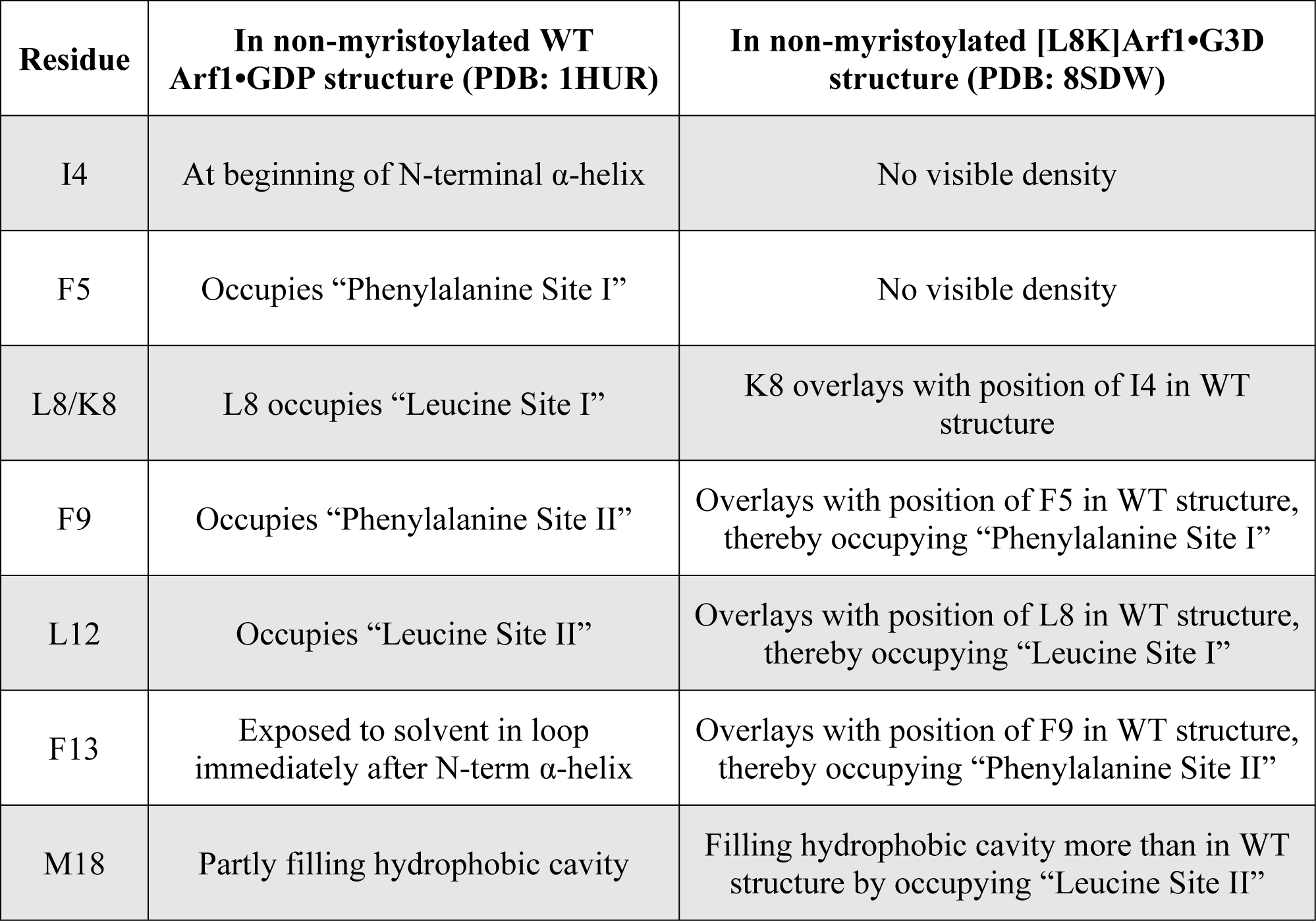
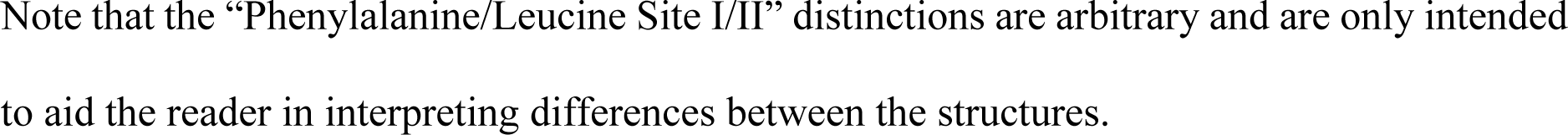
Relative positions of N-terminal HVR residues within the non-myristoylated WT Arf1•GDP and [L8K]Arf1•G3D crystal structures.

| **Residue** | **In non-myristoylated WT Arf1•GDP structure (PDB: 1HUR)** | **In non-myristoylated [L8K]Arf1•G3D structure (PDB: 8SDW)** |
| --- | --- | --- |
| I4 | At beginning of N-terminal α-helix | No visible density |
| F5 | Occupies “Phenylalanine Site I” | No visible density |
| L8/K8 | L8 occupies “Leucine Site I” | K8 overlays with position of I4 in WT structure |
| F9 | Occupies “Phenylalanine Site II” | Overlays with position of F5 in WT structure, thereby occupying “Phenylalanine Site I” |
| L12 | Occupies “Leucine Site II” | Overlays with position of L8 in WT structure, thereby occupying “Leucine Site I” |
| F13 | Exposed to solvent in loop immediately after N-term α-helix | Overlays with position of F9 in WT structure, thereby occupying “Phenylalanine Site II” |
| M18 | Partly filling hydrophobic cavity | Filling hydrophobic cavity more than in WT structure by occupying “Leucine Site II” |

Note that the “Phenylalanine/Leucine Site I/II” distinctions are arbitrary and are only intended to aid the reader in interpreting differences between the structures.

### Mutations in the N-terminal region of Arf1 may have differing effects dependent on whether Arf1 is myristoylated

The observation that the L8K mutation caused an unexpected change in non-myristoylated Arf1•GDP tertiary structure raises the question as to whether such changes might account for previously reported effects of N-terminal Arf1 mutations and how the effects might be influenced by the myristoyl moiety. [L8K]Arf1 behaved similarly to WT non-myristoylated Arf1 and WT myrArf1 in GAP assays, while a valid comparison, i.e., [L8K]myrArf1 to myrArf1, has not been examined. L8 contributes to membrane association as seen in the yeast myrArf1•GTP NMR structure (Fig. 3A); interestingly, L8 also forms hydrophobic interactions with the distal end of the myristoyl moiety in this structure (Fig. 3B). Here, we also examine the effect of mutating F13, which is surface-exposed in WT human non-myristoylated Arf1, is buried into the hydrophobic cavity in [L8K]Arf1, and is in a disordered loop exposed to solvent in yeast myrArf1 (Fig. 2B – 2D). Furthermore, F13, like L8, is one of the residues localized adjacent to the membrane in the NMR structure of *S. cerevisiae* myrArf1•GTP bound to bicelles (Fig. 3A), suggesting that this residue likely contributes to membrane association. Previous work examining the F13A mutation in non-myristoylated Arf1 found little effect on either spontaneous nucleotide exchange or recognition by Arf GAPs (GEF activity was not examined) (14). Furthermore, both these residues are among the most well-conserved in the Arf N-terminal HVRs, with only Arf6 and Arf3 exhibiting different residues at these positions (Fig. 3C). Given that these mutations (F13A and L8K) have both been previously studied in the context of non-myristoylated Arf1 and there is now evidence that one of them—L8K—causes unintended tertiary structure changes in non-myristoylated Arf1 (Fig. 1), we reexamined the effects of these mutations in the context of myristoylated protein.

**Figure 3.**
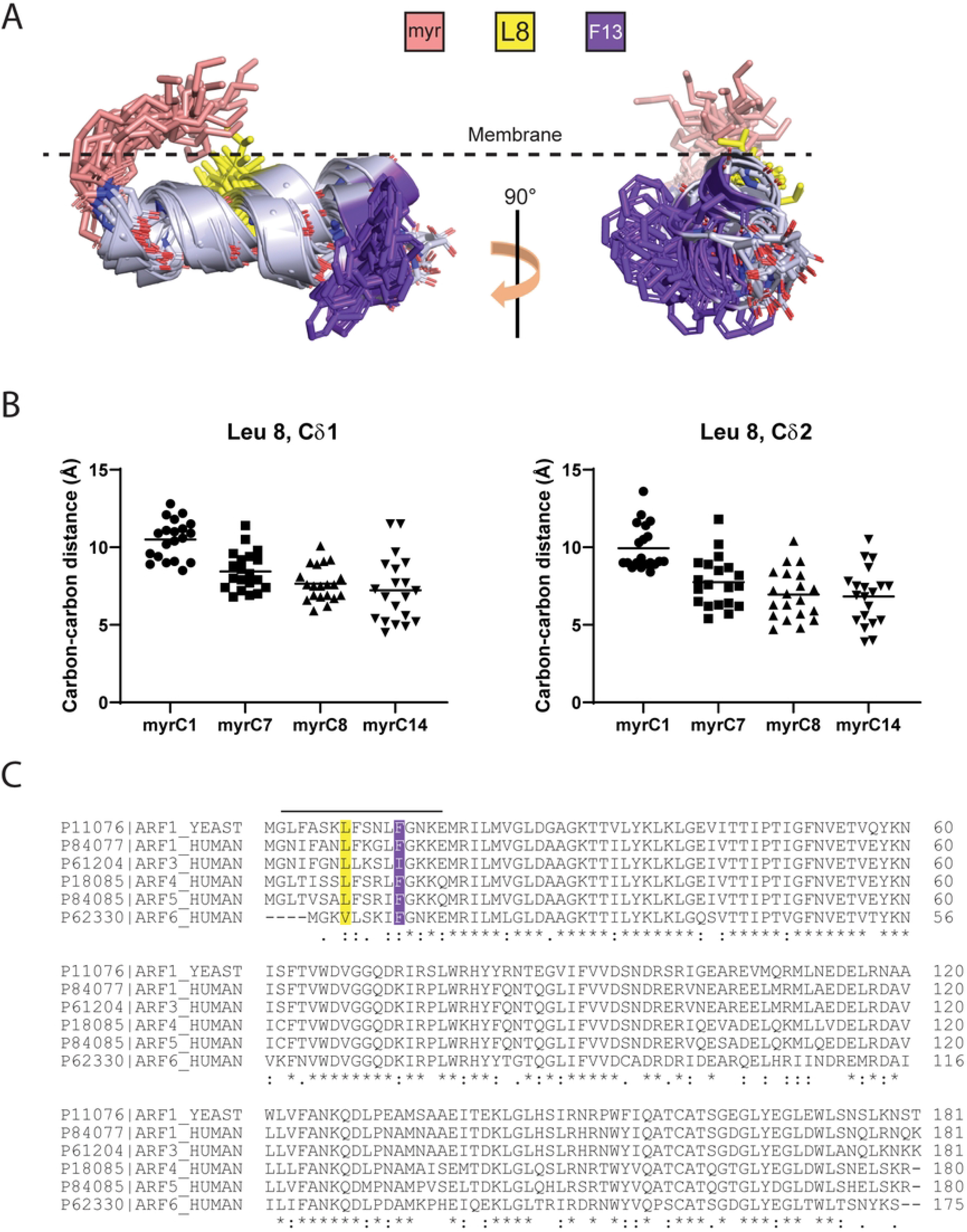
(**A**) NMR structure of the N-terminal extension of *S. cerevisiae* WT myrArf1•GTP bound to bicelles (PDB: 2KSQ (17)). Note that all NMR states of the structure are depicted. The rough positioning of the outer layer of the phospholipid membrane is shown as a dashed line, and the myristate moiety, L8, and F13 are shown as colored sticks (key on top). (**B**) Carbon-carbon distances between the indicated carbons from the myristoyl moiety and L8 in yeast WT myrArf1•GTP. Distances were obtained from each of the 20 states within the NMR structure. (**C**) Alignments of the primary sequences of *S. cerevisiae* Arf1 as well as human Arf1, Arf3, Arf4, Arf5, and Arf6. Amino acids that are identical are marked with an asterisk, whereas those with low and high similarity are marked with a period or a colon. Colors of amino acids that align with yeast L8 and F13 are as the same as in (A). The N-terminal HVRs are represented by the horizontal line on top. Swiss-Prot identifiers are shown on the left, and amino acid positions on the right.

### N-terminal mutations in myristoylated Arf1 affect neither spontaneous exchange nor recognition by the Arf GEF Brag2

The N-terminal extension of Arf undergoes a positional and conformational change on exchange of nucleotide and has an established role in nucleotide exchange (21). The contribution of individual residues within the N-terminus, however, has not been examined, particularly in the context of myristoylated Arf. Here, we examined the effects of L8 and F13 on spontaneous exchange, which was accelerated by buffering Mg^2+^ to 1 – 10 μM using EDTA. Specifically, we examined the Arf1 mutants L8K, L8A, F13A, as well as the double mutant L8A/F13A. Large unilamellar vesicles (LUVs) were included in the assay to accommodate the myristate. Compared with WT myrArf1, the rate and extent of binding GTPγS were within two-fold of one another, which, given the errors, were not significantly different (Supplemental Fig. S2). We concluded that L8 and F13 do not significantly affect spontaneous exchange in myrArf1.

Similarly, little or no change in GEF-catalyzed exchange among the mutants was detected. In our experiments, Brag2, a phosphoinositide-dependent GEF (35), was used as a model GEF. The Sec7-PH domain tandem of Brag2 (Fig. 4A, top) was titrated into a reaction mixture containing [^35^S]GTPγS, the indicated myrArf1 mutant, and LUVs containing phosphatidylinositol 4,5-bisphosphate (PIP2). The amount of Brag2 necessary for 50% of maximum observed nucleotide exchange (C_50_) was determined, which is inversely related to enzymatic power. The C_50_ of Brag2 for WT myrArf1 was 1.9 nM, and all the mutants were within ∼2-fold of that value (Fig. 4, Table 3). The C_50_ using [F13A]myrArf1 was about 2-fold greater than for WT myrArf1 or myrArf1 with mutations in L8, (p < 0.01, Fig. 4, Table 3). Thus, despite the previously documented changes in the N-terminal region of Arf1 on switching between GTP- and GDP-bound forms of Arf, neither L8 nor F13 has a dominant effect.

**Figure 4.**
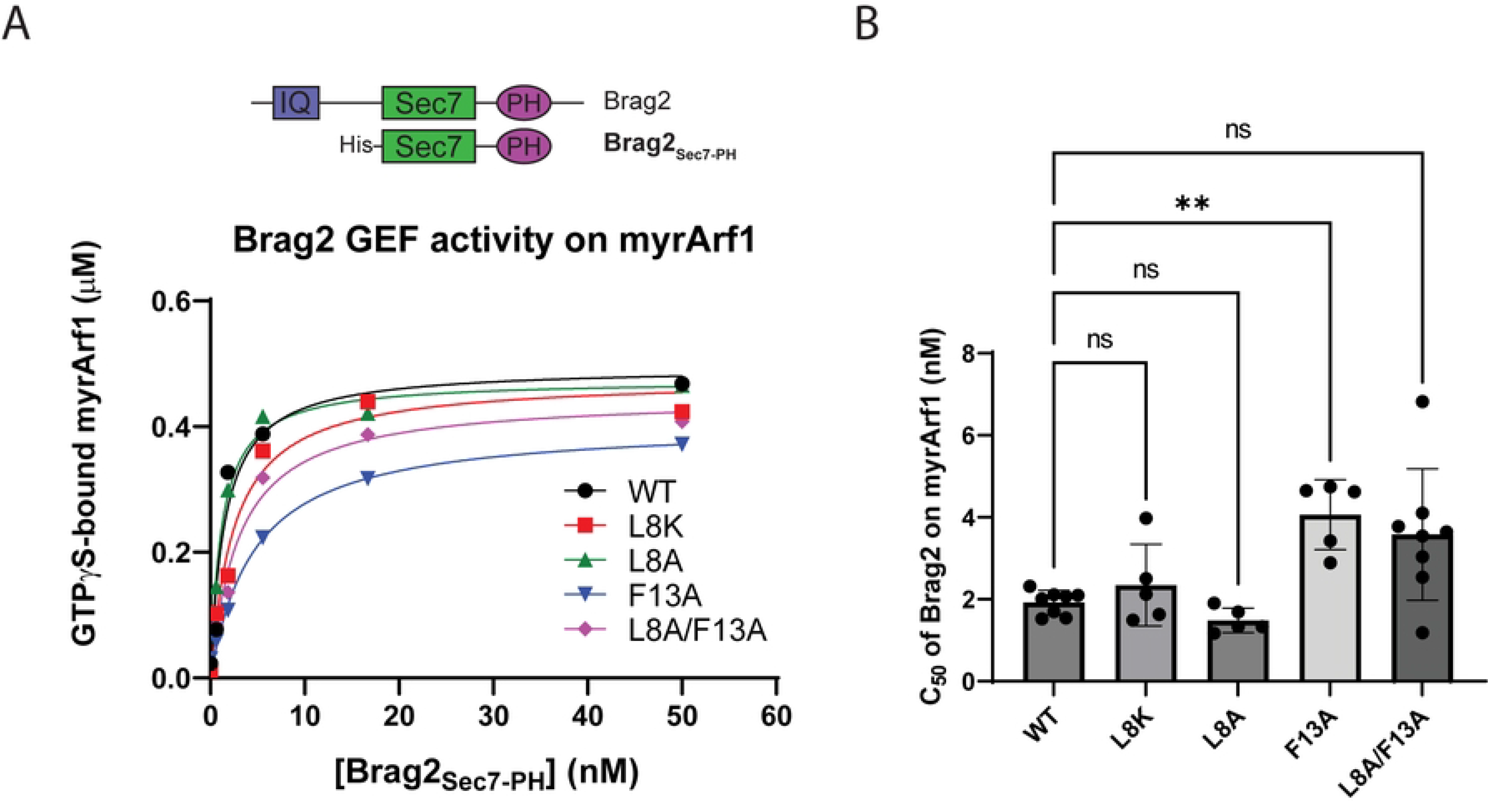
(**A**) Brag2 GEF activity using the indicated myrArf1•GDP as substrate. For these assays, the His-tagged Brag2_Sec7-PH_ construct (see top for comparison with full-length construct) was titrated into a reaction with [^35^S]GTPγS, LUVs, and 0.5 µM myrArf1 constructs. After a fixed period of time, the fraction of myrArf1 bound to [^35^S]GTPγS was measured. Data shown are a representative example from multiple experiments. IQ, IQ motif; Sec7, Sec7 catalytic domain; PH, Pleckstrin Homology domain. (**B**) Summary of Brag2 GEF activity assays using the indicated myrArf1•GDP as substrate. C_50_ values (the concentration of Brag2_Sec7-PH_ required to achieve 50% of maximum loading of the myrArf1 with GTPγS) from each independent experiment are shown. Error bars represent standard deviation. ns, not significant; **, p < 0.01 via one-way ANOVA with repeated measures (and mixed effects) and Dunnett’s multiple comparisons test against WT.

**Table 3.**
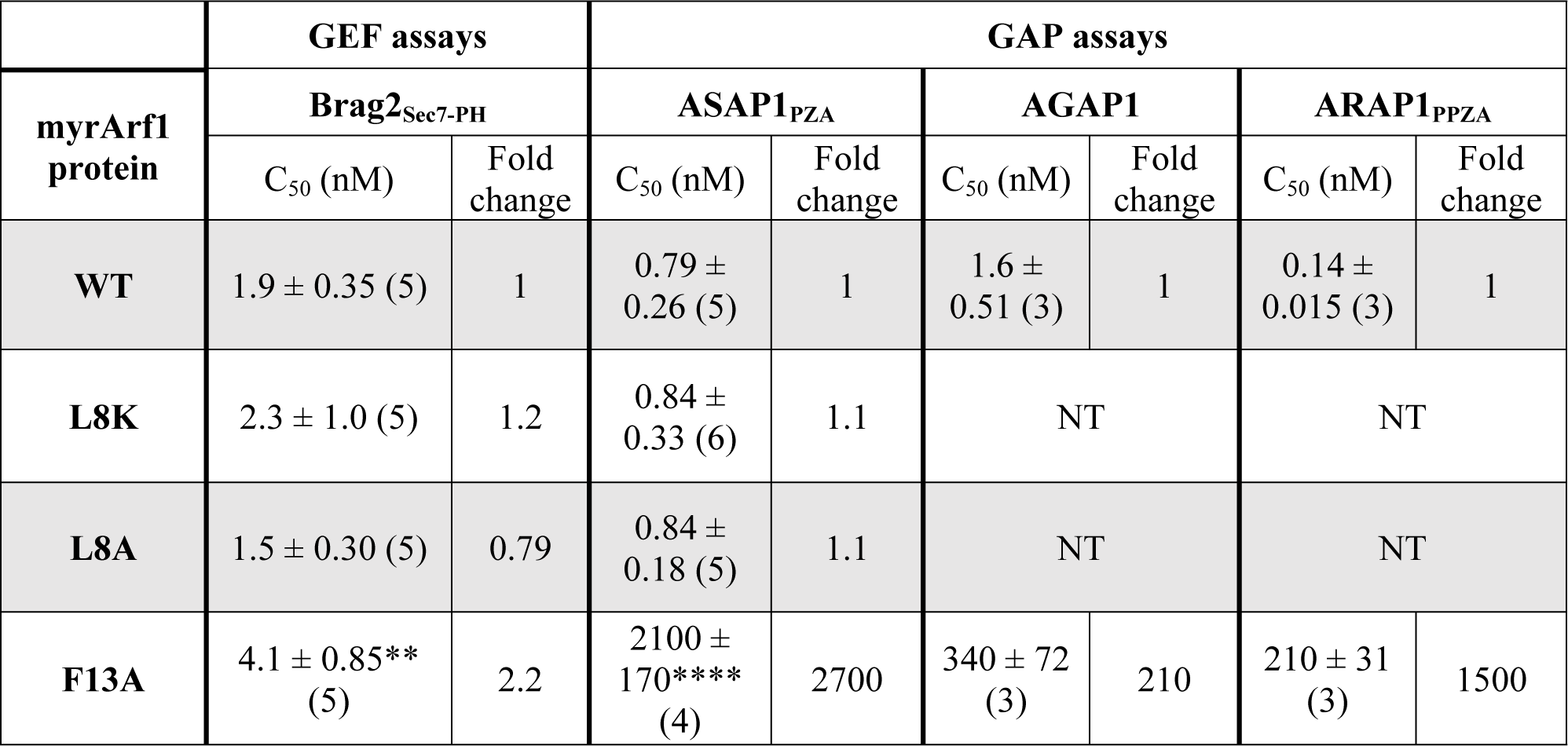
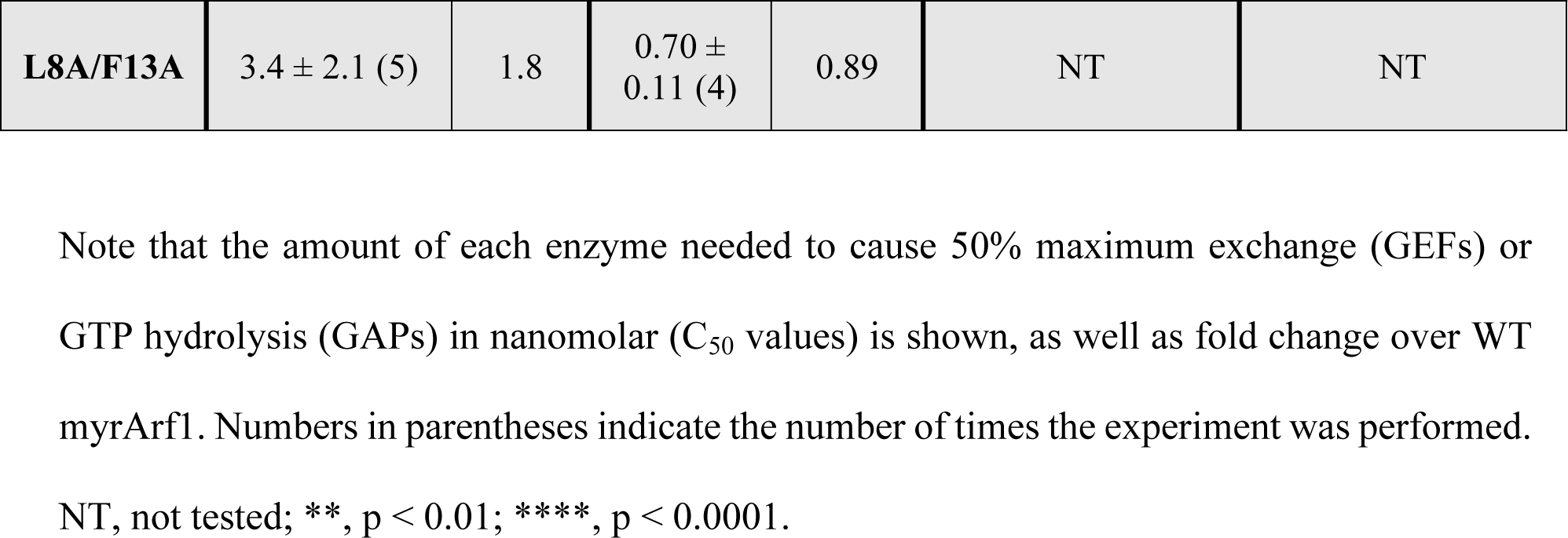
Summary of Arf GEF and GAP assays using myrArf1 constructs.

### N-terminal mutations in myrArf1 affect activity with Arf GAPs

We have previously reported a mutational analysis of the effect of the N-terminus on interaction with Arf GAPs (14). In those studies, neither L8 nor F13 were found to have a significant effect; however, they were examined in the context of non-myristoylated Arf1. Because of the possibility that the presence of the myristate could affect the N-terminus through interaction with L8 (Fig. 3), we tested each of the mutants as a substrate for the catalytic fragment of the model Arf GAP ASAP1 (Fig. 5A, 5B). ASAP1 was titrated into a reaction mixture containing the myrArf1 proteins bound to [α^32^P]GTP and LUVs containing the activating phosphoinositide PIP2. Reactions were stopped after a fixed time and the conversion of [α^32^P]GTP to [α^32^P]GDP was measured. The amount of GAP required to achieve 50% hydrolysis in the fixed time (C_50_) was estimated and, similar to the GEF reactions, is inversely proportional to enzymatic power.

**Figure 5.**
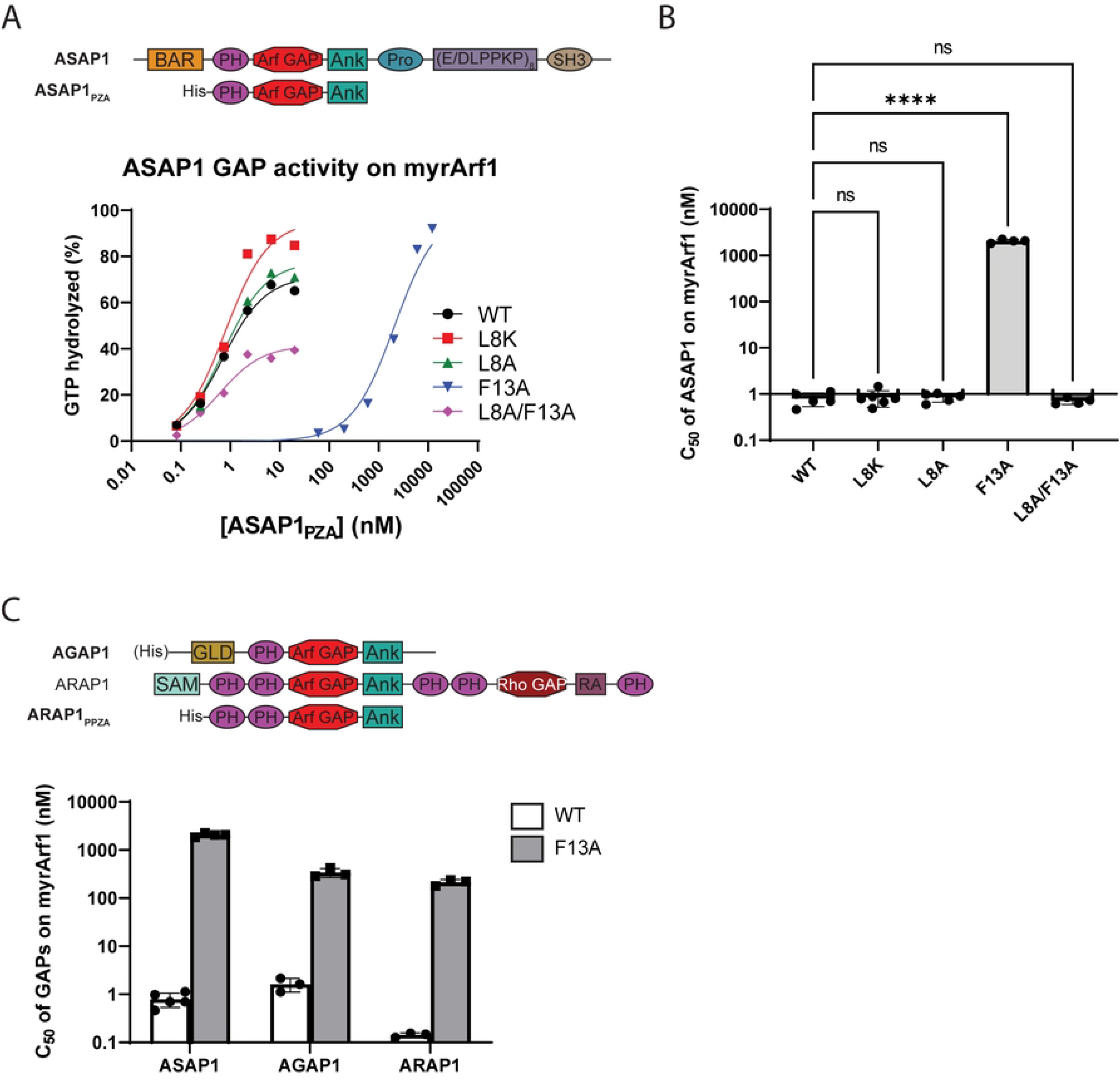
(**A**) ASAP1 GAP activity using the indicated myrArf1•GTP as substrate. For these assays, the His-tagged ASAP1_PZA_ construct (see top for comparison with full-length construct) was titrated into a reaction with myrArf1 constructs bound to [α^32^P]GTP on an LUV surface. After a fixed period of time, the ratio of [α^32^P]GDP and [α^32^P]GTP bound to myrArf1 was measured. Data shown are a representative example from multiple experiments. BAR, Bin/Amphiphysin/Rvs domain; PH, Pleckstrin Homology domain; Arf GAP, Arf GAP catalytic domain; Ank, Ankyrin repeats; Pro, Proline rich region; E/DLPPKP, E/DLPPKP repeat region; SH3, Src Homology 3 domain. (**B**) Summary of ASAP1 GAP activity assays using the indicated myrArf1•GTP as substrate. C_50_ values (the concentration of ASAP1_PZA_ required to achieve 50% of maximum GTP hydrolysis) from each independent experiment are shown. Error bars represent standard deviation. ns, not significant; ****, p < 0.0001 as determined via ordinary one-way ANOVA with Dunnett’s multiple comparisons test against WT. (**C**) Summary of GAP activity assays of His-tagged ASAP1_PZA_, full-length AGAP1, and ARAP1_PPZA_ using WT or [F13A]myrArf1•GTP as substrates. Assays were conducted as described in (A). Error bars represent standard deviation. GLD, GTP-binding protein-like domain; SAM, sterile-α motif; Rho GAP, Rho GAP catalytic domain; RA, Ras-associating domain. All other protein regions are as described in (A).

Mutation of L8 alone to either alanine or lysine had little effect on ASAP1 GAP activity, while the F13A mutation increased the C_50_ ∼2700-fold over WT myrArf1 (Fig. 5A, 5B, Table 3). The effect of the F13A mutation on the C_50_ was reversed by simultaneously including the L8A mutation, although the maximum fraction of GTP hydrolyzed was diminished compared to WT myrArf1 (Fig. 5A, 5B, Table 3; see also Supporting Information, Supplemental Fig. S3, and relevant citation (36)). To extend our analyses, we examined [F13A]myrArf1as a substrate for two additional Arf GAP subtypes, represented by AGAP1 and ARAP1 (Fig. 5C, top). For these reactions, the activating phospholipids were either PIP2 (AGAP1) or phosphatidylinositol 3,4,5-trisphosphate (PIP3, for ARAP1) (37, 38). The C_50_ values of AGAP1 and ARAP1 using [F13A]myrArf1 as a substrate were >200- and ∼1500-fold greater than for WT myrArf1 (Fig. 5C, Table 3). Given that non-myristoylated [F13A]Arf1 was previously observed to have little or no effect on ASAP1 C_50_ values compared to non-myristoylated WT Arf1 (14), the results reveal the importance of the myristate in determining function of the N-terminus.

### Molecular dynamics simulations suggest that a change of phenylalanine 13 to alanine does not affect interaction with the lipid bilayer

We considered the hypothesis that mutations in the N-terminal peptide alter its interaction with the membrane. We performed μs long molecular dynamic (MD) simulations of the wild type myristoylated N-terminal region (residues 2 – 17) and compared results to peptides carrying single point (L8A, L8K, F13A) or double point (L8A/F13A) mutations. Initially, peptides with residues 2 – 13 in alpha helical form and residues 14 – 17 in unstructured form were placed in the membrane following previous observations made by our group (Zhang *et al., Nat. Commun*, in press). Except for the L8K mutant, no helical kink or bending of the peptide was observed for the WT peptide and its alanine mutants. The peptides were located on average 6.5 ± 3.5 Å below the phosphate plane during the remainder of the simulation, with the long axis of the alpha helical segment roughly perpendicular to the bilayer normal (Fig. 6A, 6B) in good agreement with results obtained for myrArf1*•*GTP at the membrane (Zhang *et al., Nat. Commun*, in press). The hydrophobic face of the peptides points toward the bilayer core, and interactions of charged residues with lipid headgroups stabilize the peptide position (Supplemental Fig. S4A). Alanine mutations did not affect the orientation and dynamics of the myristoyl acyl chain which extends into the lipid matrix, only forming transient contacts with the peptide. On the contrary, introduction of a lysine residue in position 8 drastically changed how peptides interact with the bilayer (Supplemental Fig. S4B). On average, the orientation and depth of insertion of L8K mutant peptides are less stable along the trajectory. In addition, out of four replicas, one peptide completely loses its alpha helical structure and relocates closer to the lipid headgroup. Overall, despite exhibiting a large shift in GAP C_50_ values, we did not detect a difference in the myristoylated peptides caused by the F13A mutation, suggesting that another, as of yet to be defined mechanism renders [F13A]myrArf1 a poor substrate for Arf GAPs.

**Figure 6.**
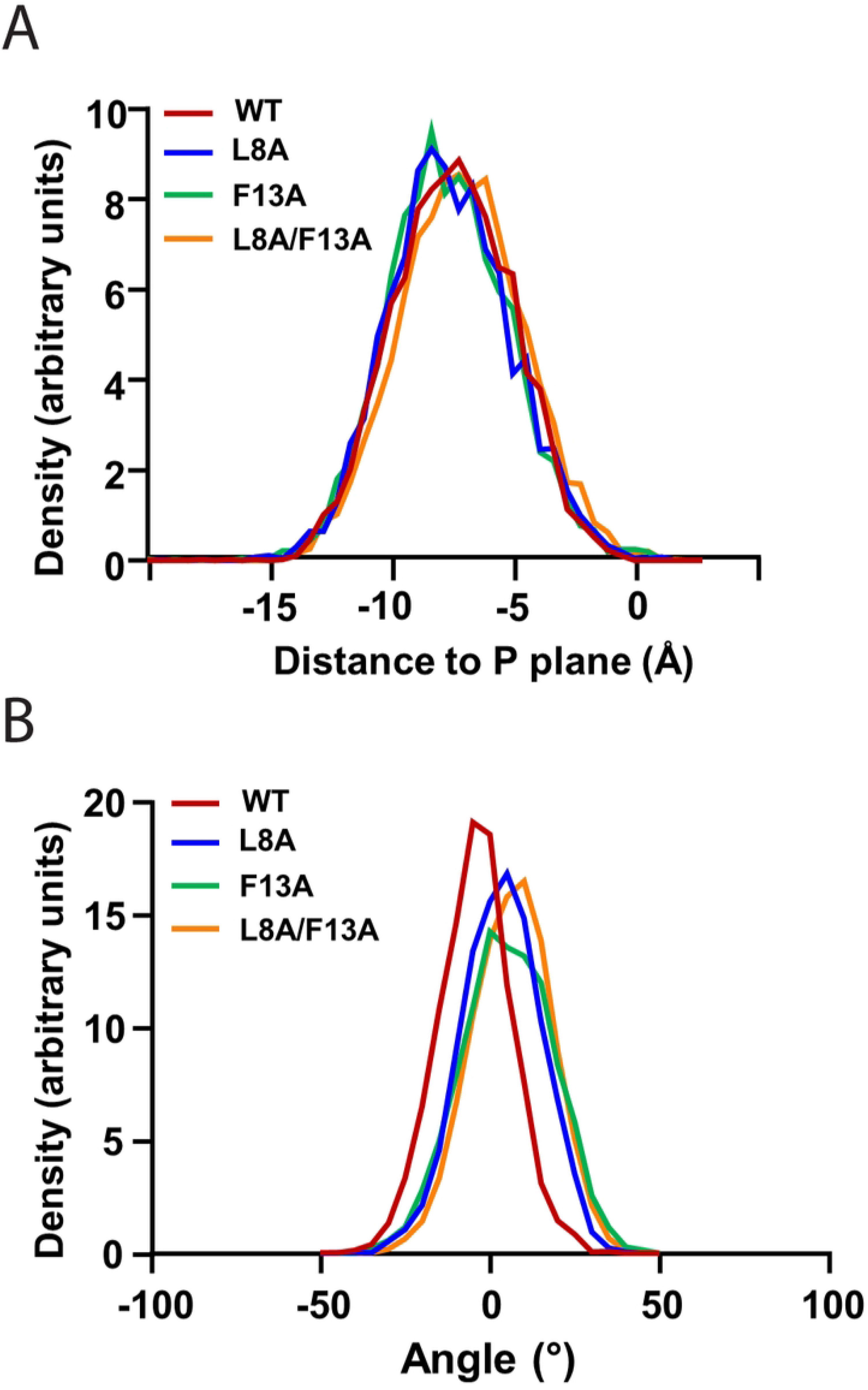
(**A**) Insertion depth of WT myrArf1 N-terminal peptide and its mutants. The location of peptides was stable along the trajectory with average depth of -6.8 ± 2.1 Å, -6.7 ± 2.2 Å, -6.9 ± 2.1 Å, and -6.1 ± 2.2 Å relative to the average location of the monolayer phosphate plane for WT, L8A, F13A, and L8A/F13A myrArf1 N-terminal peptides. (B) Average orientation of WT myrArf1 N-terminal peptide and its mutants with respect to the membrane plane. A tilt angle of zero means that the helical axis is parallel to the membrane surface. A negative tilt angle means the peptide is tilted such that the N-terminus is lower than the C-terminus on the z-axis (membrane normal).

## Discussion

Ras superfamily GTPases are commonly modified in HVRs with lipid moieties, which anchor the proteins to endomembranes. Arf GTPases, part of a subfamily of the Ras superfamily, have an N-terminal extension from the G domain that is cotranslationally and irreversibly myrsitoylated (6, 39). The myristate acts, at a minimum, as a membrane anchor; however, the potential function of the myristoylated HVR in protein-protein interactions, the molecular bases for its effects on GAP- and GEF-catalyzed transitions, and interdependence of the myristate with other functions of the HVR are understudied. Here, by examining the effect of mutating two hydrophobic residues within the N-terminal region of myrArf1, we discovered a codependence of GAP activity on the myristate and N-terminal HVR, yielding insights into the mechanism of Arf GAP activity.

### The myristate is an integral part of Arf proteins affecting the structure of the N-terminal HVR

The myristate in Arf proteins is distinguished from the lipid modifications of other Ras superfamily in that it is part of the switch, i.e., structural change, that occurs when the protein exchanges GDP for GTP (7). A comparison of the GDP- and GTP-bound yeast myrArf1 structures provides a clear illustration. In yeast myrArf1•GDP, the myristate lies in a hydrophobic cavity in the G domain upon which the N-terminal HVR floats as an unstructured peptide (23); in the GTP-bound state, the cavity is vacated, with the myristate embedding in the membrane and the HVR gaining secondary structure as an alpha helix embedded in the lipid bilayer (17). The structural role of the myristate is further illustrated by comparison of the crystal structure of human full-length non-myristoylated Arf1•GDP, in which the N-terminal region folded into an amphipathic helix that occluded the hydrophobic cavity, with the residues on the hydrophobic face inserted into the cavity (22). The myristate, therefore, is not only embedded in the GDP-bound protein but is also important to the structure of the N-terminal HVR in this state. The crystal structure of non-myristoylated [L8K]Arf1•G3D emphasizes the intrinsic nature of the myristoyl moiety for Arf proteins; the absence of the myristate in this structure, coupled with the L8K mutation, caused the N-terminal HVR to fold into an alpha helix with an unanticipated shift. Together, these structures and others provide insights into how Arf proteins are influenced by seemingly subtle changes in their N-terminal HVRs and aid our understanding of how Arf GEF and GAP catalytic mechanisms.

By comparing mutants in the context of myristoylated Arf1 with the same mutants in non-myristoylated Arf1, we gained some insight into GAP-catalyzed conversion of Arf•GTP to Arf•GDP that might involve a cooperative effect between the N-terminal HVR and the myristate. We previously reported that the N-terminal extensions of Arf proteins are required for efficient catalytic activity of PH domain-dependent Arf GAPs (12, 13). Studied in ASAP1, the mechanism entails direct binding of the N-terminus of Arf to the PH domain of the GAP. Mutational analysis in the background of non-myristoylated Arf1 identified a number of critical residues within the N-terminus for Arf GAP-catalyzed GTP hydrolysis; however, little or no effect of mutating L8 or F13 was found in the context of the non-myristoylated protein (14). In contrast, mutating F13 to alanine decreased activity almost 3000-fold when using myristoylated Arf1. By itself, mutations of L8 did not affect activity, but when mutated to alanine, reversed the effect of mutating F13. One plausible explanation of these results was that the two residues operate with the myristate to regulate alpha helical content of the N-terminal HVR, presuming Arf GAPs bind the Arf N-terminus while folded as an alpha helix. However, using MD, we did not detect any effect of the mutations on secondary structure. An alternate explanation, which we are now exploring, is related to the extraction of the N-terminal HVR from the membrane to bind the PH domain of the Arf GAP, a necessary step for catalysis. If L8 cooperates with myristate to anchor the N-terminus in the membrane and F13 makes contact with the GAP to extract the peptide from the membrane, one might expect mutation of F13 to reduce GAP activity consequent to an inability to extract the HVR from the membrane. Further, mutation of L8 to alanine would reduce the energy required to extract the peptide, thereby reversing the effect of mutating F13. We are currently developing some methods that will enable us to test predictions of this hypothesis.

Spontaneous and Brag2-catalyzed myrArf1 binding to GTP did not differ between WT and the mutants we examined. In hindsight, the lack of effect of mutating the N-terminal region might be expected, given that the N-terminal region of Arf is disordered when the myristoyl moiety is present (17). In addition, in the crystal structure of [Δ17]Arf1•G3D in complex with Brag2_Sec7-PH_ (PDB accession code: 4C0A), the N-terminal region is not localized near Brag2 (21). Thus, our results are consistent with the prevailing hypothesis that rearrangement of the interswitch domain and accommodation of the myristate within a bilayer drive the change in the N-terminal region on exchange of GDP for GTP. The N-terminal amino acids do not have a direct role for GEF-catalyzed nucleotide exchange other than facilitating interaction with the membrane via hydrophobic residues.

Given the importance of the myristoylated N-terminal HVR for interaction with GAPs (12), it might also be involved in binding effectors, which is understudied. Most structures of Arf•GTP in complex with effectors have been solved using truncated Arf proteins (40). A unifying feature among the effector:Arf•GTP interface regions is a so-called “common hydrophobic area” that is ∼480 Å^2^ and comprised of residues within switch I, interswitch, and switch II regions of the G domain. This common hydrophobic area acts as a scaffold upon which effector proteins with diverse functions bind to the Arf G domain, the latter itself being considered quite rigid (40). However, as many of these structures were not solved with a membrane mimetic, nor the native (myristoylated, full-length) protein, it is possible that the HVRs may also contribute to Arf:effector interactions in addition to the common hydrophobic area, which would not be captured by these structures. A compelling example of how these interactions may be lost in these structures can be seen by examination of the co-crystal of full-length Arl2•GTP in complex with the effector Binder of ARL Two (BART). In addition to interactions with switch I in Arl2, BART also made extensive hydrophobic interactions with the N-terminal region of Arl2, as the Arl2 N-terminal alpha helix was found to reside within a hydrophobic cleft on the BART surface (41). The myristoylated N-terminal HVRs, which are the most different among the Arf proteins, might therefore provide specificity for the association with effectors in some circumstances. Another possible mechanism for controlling protein-protein interactions is orientation. For example, the N-terminal extensions of Arf1 and Arf6 are 16 and 12 amino acids in length; given that alpha helices exhibit a characteristic 3.6 residues per helical turn (42), the packing of the helices at the C-terminal end of the HVRs may be different, which could in principle constrain the three-dimensional space available for the G domain to sample within the cytosol.

Our results raise the question of how the properties of Arf proteins might extend to other Ras superfamily members. The HVR has been largely ignored, with much focus instead being centered on the G domain. Considered just a membrane anchor (43), some *in vitro* studies will replace the HVR with a His tag to Ras proteins of interest and use a lipid derivatized with a Nickel-NTA moiety to anchor the protein to a membrane (44). Even in cases where the HVR clearly determines specificity, inference has been made using the truncated Ras protein. For example, Sin1, a subunit of the target of rapamycin complex 2 (TORC2) specifically binds Ras4A dependent on the Ras4A HVR, but the crystal structure of a mutant form of Ras4B lacking the HVR was instead used to solve the co-crystal structure in complex with Sin1 (45). Furthermore, the specific lipid modification is also important for some Ras interactions—in phosphodiesterase (PDE) δ, the interaction with Ras is mediated specifically by the farnesyl group and a stretch of six amino acids from the HVR (46). It is therefore likely that exclusion of the acylated HVRs of Ras leads to an oversimplification of the system, and as such, that these features (and membrane mimetics) should be included whenever possible in order to best represent the behavior of these proteins *in vivo*.

In summary, we provide a crystal structure of an Arf1 variant, and also observed a cooperative effect of the myristate and N-terminal HVR that controls Arf GAP activity. These results provide insights into the Arf GAP catalytic mechanism. Further, we highlight the value of utilizing GTPases in their native form (or as close as possible) for *in vitro* studies designed to determine the mechanism of action of various Arf regulators or effectors.

## Materials and Methods

### Chemicals

Both [α^32^P]GTP and [^35^S]GTPγS were obtained from Perkin-Elmer.

### Protein expression and purification

Non-myristoylated [L8K]Arf1 (8), the Sec7 and PH domains (Sec7-PH construct) of Brag2 (both His-tagged for assays, and GST-tagged for myrArf1 purifications, see below) (35), the PH, Arf GAP, and ankyrin repeats (PZA construct) of ASAP1 (47), full-length AGAP1 (48), and the tandem PH domains, Arf GAP, and ankyrin repeats of ARAP1 (PPZA construct) (38) were expressed and purified as previously described.

Expression of myrArf1 proteins with C-terminal 6x His tags was accomplished via co-expression with yeast N-myristoyltransferase (yNMT) using the pETDuet-1 vector (Novagen) in *E. coli* BL-21(DE3) as previously described (49). Expressed myrArf1 proteins were purified from the insoluble fraction (GTP-bound protein) following lysis and ultracentrifugation in 20 mM Tris pH 8.0, 100 mM NaCl, 1 mM MgCl_2_, and 1 mM DTT (TNMD buffer) supplemented with cOmplete EDTA-free protease inhibitor cocktail (Roche). The insoluble myrArf1•GTP was solubilized using 20 mM Tris pH 8.0, 100 mM NaCl, and 1 mM MgCl_2_ (TNM buffer) supplemented with 1% Triton X-100, and ultracentrifuged to remove remaining insoluble material. Solubilized myrArf1•GTP was enriched via ammonium sulfate precipitation (35% at 0°C) and ultracentrifugation, whereupon it salted out as a membrane film. The supernatant was removed, and the membrane was resuspended in TNM buffer + 1% Triton X-100. The GTP-bound myrArf1 was then converted to its GDP-bound state by addition of EDTA to 2 mM (to buffer Mg^2+^ concentrations to 1 – 10 µM), GDP to 10 mM, and GST-tagged Brag2_Sec7-PH_ to ∼0.6 µM, and subsequent incubation for 1 – 2 hours at 30°C. After, the exchange reaction was quenched by addition of MgCl_2_ to 2 mM. The sample was then diluted fivefold with fresh TNM, ultracentrifuged to clarify, and the soluble GDP-bound protein was loaded onto a 1 mL HisTrap HP nickel column (Cytiva) pre-equilibrated with 20 mM Tris pH 8.0, 500 mM NaCl, 20 mM imidazole (His buffer A). The myrArf1•GDP protein was washed with ≥10 column volumes (CV) of His buffer A to remove Triton X-100 and GST-Brag2_Sec7-PH_, and was then eluted using a linear gradient of 10 CV to His buffer B (20 mM Tris pH 8.0, 500 mM NaCl, 500 mM imidazole). Eluates containing myrArf1•GDP were further purified by size exclusion chromatography (SEC) using a HiLoad 26/600 Superdex 75 prep grade column (GE Healthcare) pre-equilibrated and then run in an isocratic manner using fresh TNMD buffer at 4°C. SEC eluates of sufficient purity were pooled, concentrated using a 10,000 MWCO concentrator (Amicon), aliquoted, and snap-frozen using liquid nitrogen.

### Crystallization of [L8K]Arf1•G3D and X-ray data collection and processing

Purified (≥95%) [L8K]Arf1 protein was concentrated to 10 mg/mL in TNMD buffer and was subsequently used to set crystal trays at 4°C. Initial screens were performed with the protein pre-incubated with small molecule ligands using a Mosquito system (TTP Labtech) and several commercial crystal screens from Hampton Research, Molecular Dimensions, and Emerald BioSystems by the hanging-drop method. The following day, ∼13% of all crystal drops exhibited needle-like crystals. A tray of the *apo* protein was then set using the Mosquito system and the PEG/Ion commercial screen (Hampton Research) using a 1:1 drop ratio of protein to precipitant (250 nL each). Multiple hits were found the following day, but one of the largest needle-like crystals was in formulation #17 from the screen (0.2 M Sodium nitrate, 20% w/v Polyethylene glycol 3,350, pH 6.8). This crystal was harvested directly from the screening tray after approximately a week and a half, without cryoprotectant, and was used for data collection.

Data collection was performed at the Advanced Photon Source (APS) synchrotron beamline 22-ID with a cryostream set to 100 K. The wavelength used for data collection was 1.0 Å, and a full data set was collected to 1.75 Å resolution using a Dectris EIGER X 16M detector (see Table 1 for details of data collection). The data sets were processed using HKL-2000 (50) and the intensity data was then imported into the Phenix software suite (51) as an .mtz file. The structure of the protein was solved using Phaser (52) to perform molecular replacement with a monomer of non-myristoylated Arf1•GDP (PDB entry 1HUR) (22). Examination of the initial molecular replacement solution in Coot (53) showed numerous obvious errors in the placements of residues within the interswitch region of the G domain, as well as the N-terminal HVR. To fix these issues, the residues in these regions were removed, and then manually built in Coot using the positive F_o_ – F_c_ electron density, contoured at 3σ, as a guide. The updated coordinates containing all visible amino acid residues was then iteratively refined in Phenix and Coot (see Table 1 for details of refinement). Due to the large variations in the L8K structure compared to our search model (a 1HUR monomer, which exhibited an R_free_ of ∼0.34 with our data), phase bias was not considered a significant concern. The final structure exhibited no Ramachandran outliers and ∼1.2% of residues were in the allowed region. Structures were visualized in PyMOL (Schrödinger) (54). RMSD calculations were made using the SuperPose web server (55).

### Cloning of Arf1 mutants

The Arf1 mutants L8K, L8A, F13A, and L8A/F13A were generated by site-directed mutagenesis of the Arf1/yNMT in pETDuet-1 plasmid described above as a template. Briefly, reactions were performed with mutagenesis primers and the Q5 DNA polymerase (New England Biolabs). Afterward, the template DNA was digested using *DpnI* enzyme (New England Biolabs). Mutagenized DNA was transformed into NEB® 5-alpha competent *E. coli* (New England Biolabs), and individual colonies were cultured, used to generate miniprepped DNA with a commercial kit (Qiagen), and sequenced with pET Upstream primer (Novagen). Sequencing was conducted at the Center for Cancer Research (CCR) Genomics Core at the National Cancer Institute, Bethesda, MD. Plasmid DNA with the correct sequences were then transformed into BL-21(DE3) *E. coli* for large-scale expression.

### Preparation of LUVs

LUVs were prepared by extrusion. Briefly, 1 µmol lipids, dissolved in chloroform in a siliconized glass tube, with molar ratio of 40% phosphatidylcholine, 25% phosphatidylethanolamine, 15% phosphatidylserine, 10% cholesterol, and 10% total phosphoinositide (2.5% phosphatidylinositol 4,5-bisphosphate [PIP2], 0.5% phosphatidylinositol (3,4,5)-trisphosphate, and 7% phosphatidylinositol [PI] for ARAP1 GAP activity, 1% PIP2 and 9% PI for all other experiments) were dried under a nitrogen stream for 30 minutes to 1 hour, followed by lyophilization for at least one hour. The dried lipids were resuspended in 200 µL 1x PBS, for a final concentration of 5 mM. The solution was vortexed, subjected to five rounds of freeze/thaw, and extruded using a lipid extruder (Avanti Polar Lipids) through a Whatman Nuclepore Track-Etched membrane with 1 µM pores. The LUVs were stored at 4°C and were used within a week for activity assays.

### Spontaneous guanine nucleotide exchange assays

Spontaneous exchange assays were performed as previously described (7, 12). Exchange reaction mixtures contained 25 mM HEPES, pH 7.4, 100 mM NaCl, 1 mM dithiothreitol, 0.5 mM MgCl_2_, 1 mM EDTA, 5 µM GTPγS spiked with [^35^S]GTPγS, 0.5 mM LUVs, and 0.5 μM myrArf1•GDP. The reactions were incubated at 30°C for indicated periods of time and then quenched with 2 mL of ice-cold 20 mM Tris, pH 8.0, 100 mM NaCl, 10 mM MgCl_2_, and 1 mM dithiothreitol. Protein-bound nucleotide was trapped on nitrocellulose, and the bound radioactivity was quantified by liquid scintillation counting.

### GEF activity assays

GEF activity assays were performed as previously described (35). Reaction mixtures contained 25 mM HEPES, pH 7.4, 100 mM NaCl, 1 mM dithiothreitol, 2 mM MgCl_2_, 1 mM EDTA, 5 µM GTPγS spiked with [^35^S]GTPγS, 0.5 mM LUVs, 0.5 μM myrArf1•GDP, and variable concentrations of Brag2_Sec7-PH_. The reactions were incubated at 30°C for 3 min. and quenched with 2 mL of ice-cold 20 mM Tris, pH 8.0, 100 mM NaCl, 10 mM MgCl_2_, and 1 mM dithiothreitol. Protein-bound nucleotide was trapped on nitrocellulose, and the bound radioactivity was quantified by liquid scintillation counting.

### GTP hydrolysis and GAP activity assays

GAP-induced conversion of myrArf1•GTP to myrArf1•GDP was determined as described previously (13, 56). Reaction mixtures contained 25 mM HEPES, pH 7.4, 100 mM NaCl, 1 mM dithiothreitol, 2 mM MgCl_2_, 1 mM GTP, 0.5 mM LUVs, myrArf1 bound to [α^32^P]GTP, and variable concentrations of Arf GAP. The LUVs were included in the myrArf1 GTP loading reaction. The reactions were incubated at 30°C for 3 minutes (unless otherwise specified), and quenched with 2 mL of ice-cold 20 mM Tris, pH 8.0, 100 mM NaCl, 10 mM MgCl2, and 1 mM dithiothreitol. Protein-bound nucleotide was trapped on nitrocellulose, and guanine nucleotide was released by addition of formic acid. [α^32^P]GDP and [α^32^P]GTP were then separated using thin-layer chromatography plates, and quantified.

### Molecular dynamics simulations

We modelled the amphipathic helix (residues 2-17) of myr-Arf1, including the N-terminal myristoyl group in a membrane bilayer. Five simulations were performed including WT myr-Arf1 and mutations L8K, L8A, F13A and L8A/F13A. The system was built in the CHARMM-GUI membrane builder. Each bilayer contained two of the amphipathic helices evenly distributed in each monolayer, 9 Å above the center of the bilayer. All equilibration and MD simulations were performed using AMBER with the CHARMM force field and solvated by TIP3P water molecules. The membrane contained 150 lipids in each monolayer with 142 (DMPC) and 8 PI(4,5)P2 lipids. The system was solvated with ∼15,000 TIP3 water molecules. Each simulation contained 40 Cl^−^ ions and 100 K^+^ ions with the exception of L8K, which included 96 K^+^ ions. The simulation box had dimensions measuring 129 Å × 129 Å × 70 Å. The system was equilibrated for six steps (steps 1-3 for 125 ps and 4-6 for 500 ps). Simulations were then performed for 1.5-2.5 µs at 303.15 K.

|  | **Total # of atoms** | **# of TIP3 water** | **# of μs** |
| --- | --- | --- | --- |
| WT | 82324 | 15056 | 2.29 |
| L8K | 82408 | 15092 | 1.78 |
| L8A | 82204 | 15028 | 1.79 |
| F13A | 82422 | 15102 | 2.49 |
| L8A/F13A | 82323 | 15081 | 2.48 |

### Orientation and depth of insertion from MD Structures

The tilt of each peptide was obtained by calculating the angle between the smallest moment of inertia of the alpha helical segment and the membrane surface. The tilt angle is equal to zero when the axis of the smallest moment of inertia is parallel to the membrane surface and negative when the N-terminus is lower than the C-terminus on the z axis (membrane normal). The depth of insertion of a peptide was calculated as the distance between the center of mass for heavy backbone atoms and the center of the bilayer.

## Acknowledgements

We would also like to thank Drs. Richard A. Kahn and David Lambright for their insights into the molecule G3D.

## Supporting Information Captions

**Supplemental Figure S1.**
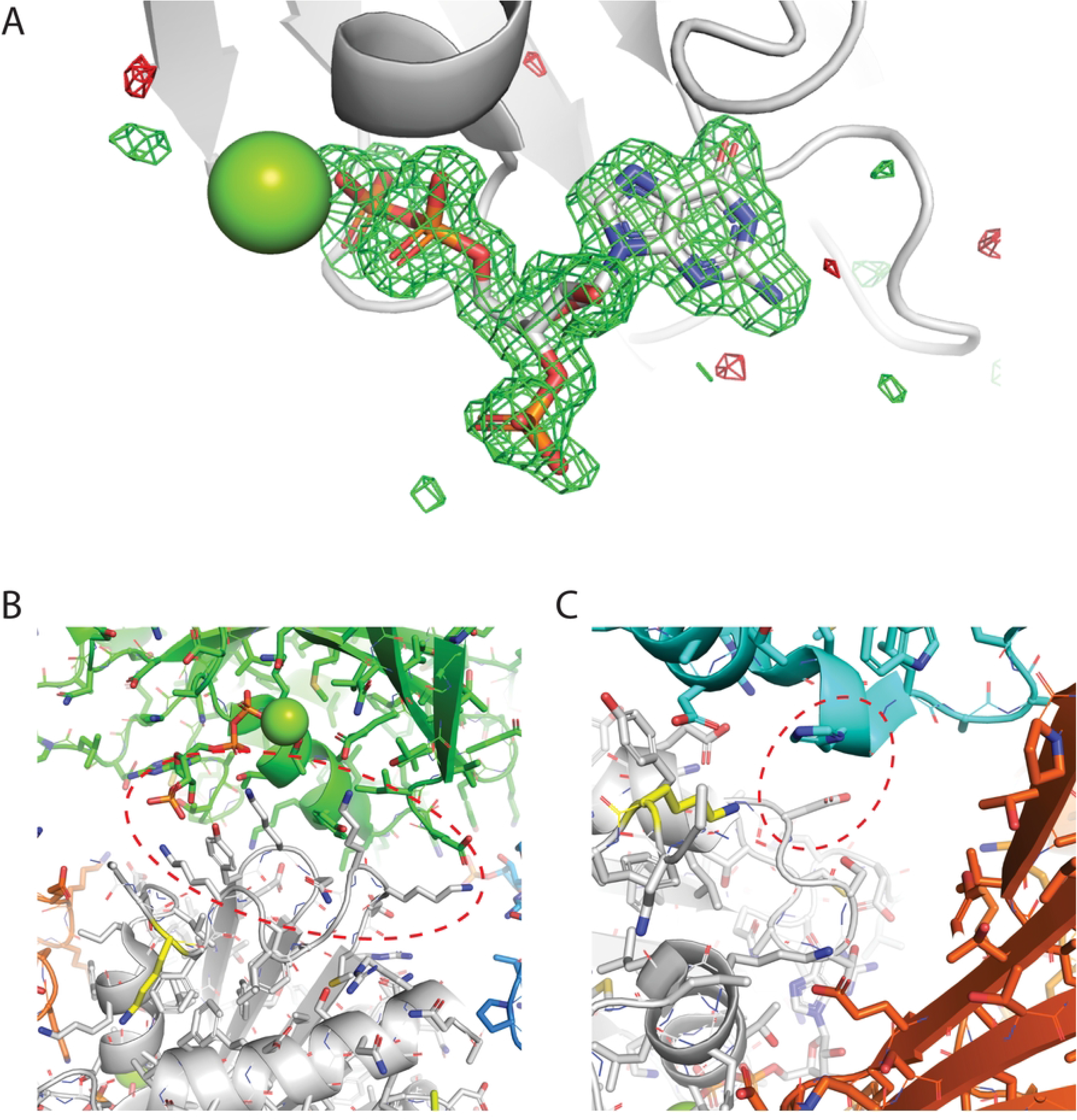
(**A**) Polder OMIT map (57) of the atoms composing guanosine-3’-monophosphate-5’-diphosphate (G3D) within the [L8K]Arf1•G3D crystal structure. Green and red mesh corresponds to positive and negative mF_obs_ − DF_model_ OMIT difference density contoured at 3σ. (**B**) Interswitch region crystal contacts between monomers within the [L8K]Arf1•G3D crystal structure. The interswitch region of one monomer (white) is adjacent to an alpha helix between the P-loop and switch I in the G domain of another monomer (green). Crystal contact regions are emphasized with red dashed line. (**C**) G5 motif crystal contacts between monomers within the [L8K]Arf1•G3D crystal structure. The G5 motif of one monomer (white) is adjacent to an alpha helix towards the C-terminal end of the G domain of another monomer (cyan). Crystal contacts are emphasized as in (B).

**Supplemental Figure S2.**
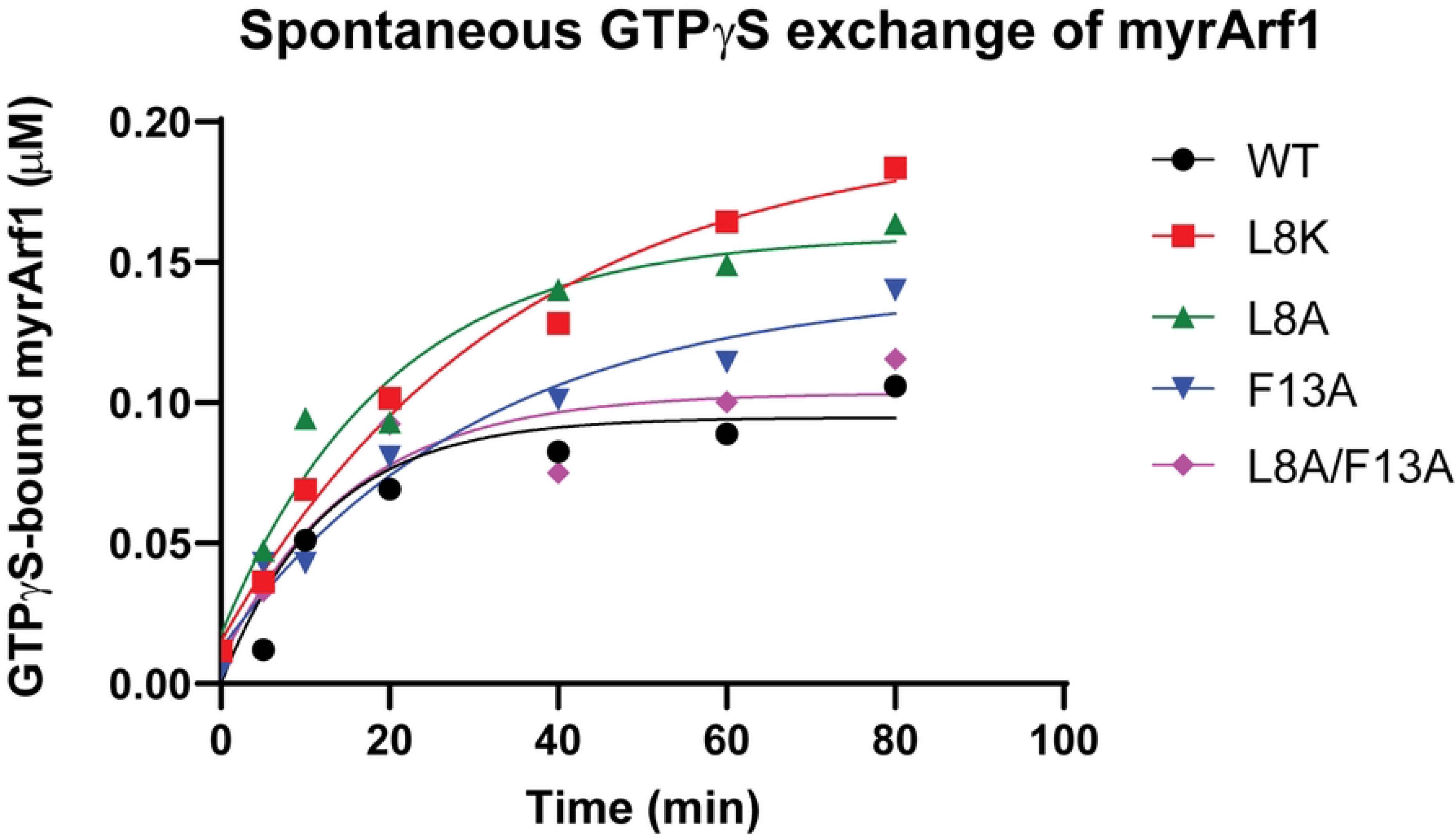
Spontaneous guanine nucleotide exchange of myrArf1 constructs. For these assays, 0.5 µM myrArf1 constructs was added to a reaction with low (1 – 10 μM) Mg^2+^ to promote exchange, as well as [^35^S]GTPγS and LUVs. After the indicated period of time, the fraction of myrArf1 bound to [^35^S]GTPγS was measured. Data shown are a representative example from multiple experiments.

**Supplemental Figure S3.**
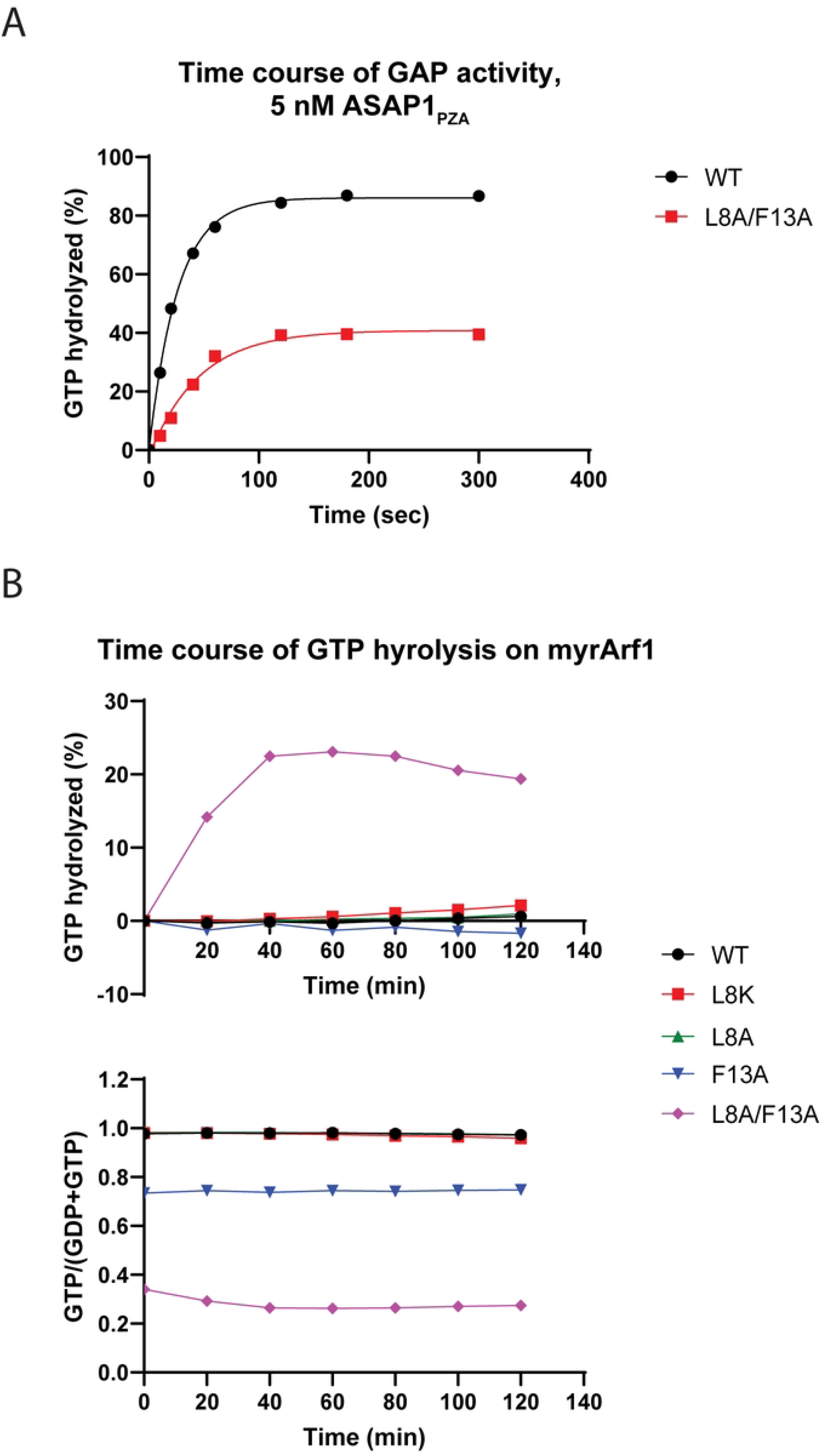
(**A**) Time course of Arf GAP activity with high concentrations (5 nM) of ASAP1_PZA_ using WT or [L8A/F13A]myrArf1•GTP as substrates. For these assays, myrArf1 was loaded with [α^32^P]GTP, ASAP1_PZA_ was added, the reaction was quenched after the indicated incubation time, and the ratio of [α^32^P]GDP and [α^32^P]GTP bound to myrArf1 was measured. Data shown are a representative example from multiple experiments. (**B**) Time course of spontaneous GTP hydrolysis on myrArf1 constructs. For these assays, myrArf1 was loaded with [α^32^P]GTP for 30 minutes. Following the indicated periods of time after addition of GAP reaction buffer containing Mg^2+^ and GTP, the reaction was quenched and the ratio of [α^32^P]GDP and [α^32^P]GTP bound to myrArf1 was measured. The panel on top shows the change in GTP hydrolyzed compared to that at time 0 minutes (immediately after 30 minutes of GTP loading then quenching); the panel on bottom is the same data, showing the full ratio of measured GTP over the sum of measured GDP and GTP. Data shown are a representative example from multiple experiments.

**Supplemental Figure S4.**
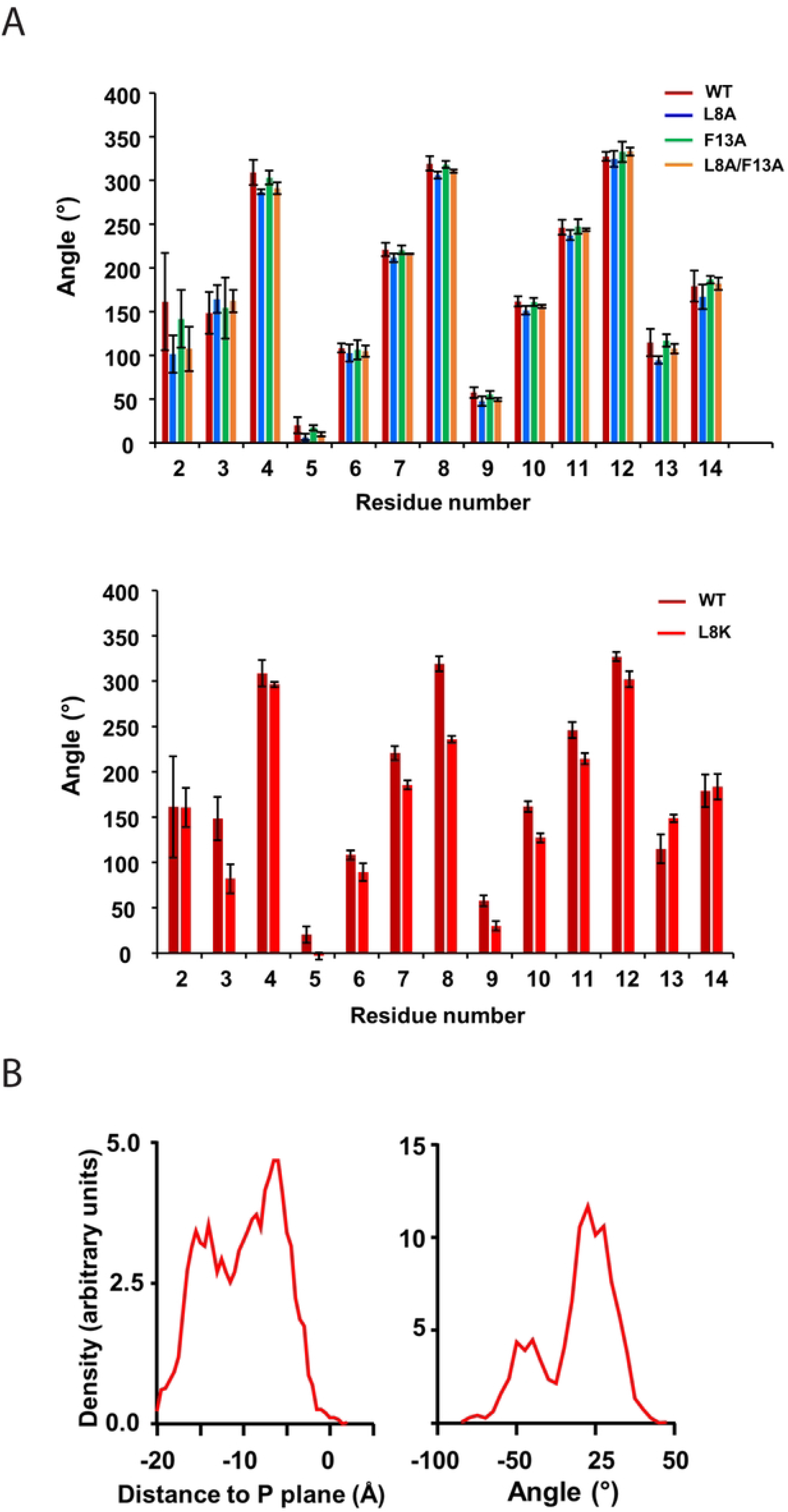
(**A**) Per residue roll angle for WT myrArf1 peptide and its mutants. WT myrArf1 per residue roll angles are compared to [L8A]myrArf1, [F13A]myrArf1, and [L8A/F13A]myrArf1 (top) or [L8K]myrArf1 (bottom). Roll angles are calculated as the angle between the C^alpha^-H^alpha^ bond of a residue and the bilayer normal. An angle of 0° corresponds to the C^alpha^-H^alpha^ bond aligned with the bilayer normal and pointing toward the hydrophobic core of the membrane. (**B**) Insertion depth (left) and average orientation (right) relative to the monolayer phosphate membrane plane of [L8K]myrArf1 N-terminal peptide. A tilt angle of zero means that the helical axis is parallel to the membrane surface. A negative tilt angle means the peptide is tilted such that the N-terminus is lower than the C-terminus on the z-axis (membrane normal).

